# Stability and DNA Methyltransferase Activity of DNMT3A are Maintained by Ubiquitin-Specific Peptidase 11 (USP11) and Sumoylation Countering Degradation

**DOI:** 10.1101/2025.03.05.641683

**Authors:** Taishi Yonezawa, Justine C. Rutter, Raghav Ramabadran, Venkatasubramaniam Sundaramurthy, Gandhar Datar, Mikolaj Slabicki, Margaret A. Goodell

## Abstract

DNA methyltransferase 3A (DNMT3A) plays crucial roles in mammalian development and hematopoiesis. DNMT3A protein instability is associated with blood diseases such as myelodysplastic syndrome (MDS), acute myeloid leukemia (AML), as well as Tatton-Brown-Rahman syndrome, an overgrowth disorder. We found that certain unstable DNMT3A mutations cause DNMT3A localization changes, resulting in loss of function. This mislocalization is partially rescued by E1 enzyme inhibition or stable USP11 expression, as DNMT3A stability is maintained by deubiquitinating enzyme USP11 countering degradation by CUL4-DCAF8 E3 ligase. We also found that USP11 enhances DNMT3A SUMOylation by promoting the interaction between DNMT3A and SUMO E3 ligases. DNMT3A SUMOylation also is essential to maintain DNMT3A protein stability. Furthermore, we found USP11 enhances the binding of DNMT3A to the polycomb complex and maintains DNMT3A DNA methyltransferase (MTase) activity. This study uncovers the mechanism for DNMT3A protein turnover through USP11, which is essential to DNMT3A function, and may be a therapeutic approach for diseases caused by DNMT3A protein instability.

## Introduction

DNA methyltransferase 3A (DNMT3A) plays an important role in the regulation of hematopoietic stem cell (HSC) differentiation [1]. Somatic DNMT3A mutations result in the clonal expansion of mutant HSCs, a phenomenon called clonal hematopoiesis (CH). Many DNMT3A mutations have been identified in CH, which accelerates with age and is associated with an ∼11-fold increase in risk of hematologic disorders like MDS and increased all-cause mortality [2, 3]. DNMT3A mutations are also associated with developmental growth disorders like Tatton-Brown-Rahman syndrome (TBRS). The symptoms of TBRS include intellectual disabilities, seizures, overgrowth, multiple developmental abnormalities, and a predisposition to cancer [4, 5]. Thus, DNMT3A is involved in hematopoiesis and development, and it is thought to be a tumor suppressor and developmental regulator.

Our recent study indicated that ∼30% of CH-related DNMT3A mutations downregulate its expression at the protein level, and some of the DNMT3A mutations associated with CH overlap with those found in TBRS [6]. The dominant-negative DNMT3A R882 hotspot mutation accounts for approximately 60% of mutations in AML [7–9] and compared to WT, the R882 hotspot mutant displays the same level of protein expression. However, this mutation accounts for only around 20% of CH cases and the mutant protein does not differ significantly in protein expression compared to wild type (WT) DNMT3A. In CH, there is a larger proportion of DNMT3A unstable variants.

Additionally, a patient cohort showed that unstable DNMT3A mutants are an independent risk factor of AML even after accounting for age and sex. The hazard ratio of unstable DNMT3A mutants is higher than the R882 and other mutations, indicating that unstable DNMT3A is strongly associated with CH and MDS risk [10]. Consequently, we hypothesized that the unstable DNMT3A mutation’s protein remark should be restored to normal for the treatment of DNMT3A-related hematologic malignancies and TBRS. Thus, it is essential to understand unstable DNMT3A mutants at the molecular, animal, and clinical levels.

Using CRISPR screening in 293T cells [11], we recently revealed that the E3 ubiquitin ligase adaptor DCAF8 induced protein degradation in unstable DNMT3A mutants [6]. Additionally, we found that WT DNMT3A is less susceptible to degradation by the ubiquitin system than unstable DNMT3A mutants. Based on these results, we hypothesize that WT DNMT3A has increased stability through deubiquitination. Furthermore, understanding the mechanisms of DNMT3A protein turnover will be useful to develop a new therapeutic approach for hematologic malignancies and TBRS. However, which deubiquitinating enzyme induces DNMT3A stabilization remains unknown.

In this study, we used a CRISPR screen in a neuroblastoma cell line to confirm the broader role of DCAF8 as a degrader of DNMT3A through ubiquitination. Through the CRISPR screening and immunoprecipitation mass spectrometry (IP-MS), we showed that Ubiquitin-Specific Peptidase 11 (USP11) induces WT DNMT3A deubiquitylation and stabilization countering degradation by DCAF8. We also found that USP11 accelerates DNMT3A SUMOylation and that this SUMOylation is essential for DNMT3A protein stabilization.

Furthermore, we found USP11 enhances the binding of DNMT3A1 (full-length) but not DNMT3A2 (short isoform) to polycomb complexes and maintains DNMT3A DNA methyltransferase (MTase) activity. DNMT3A1 and polycomb complexes are essential for neuronal gene expression and mouse postnatal development [12, 13].

Collectively, this study shows that DNMT3A protein turnover is maintained by the deubiquitinating enzyme USP11, and SUMOylation of DNMT3A by USP11 modulates DNMT3A DNA Mtase activity. Thus, regulating the activity of USP11 may be a new therapeutic strategy for disorders caused by unstable DNMT3A.

## Results

### CRISPR screening identified DNMT3A protein modulators

To better understand the mechanisms of DNMT3A protein turnover, we first confirmed DNMT3A and DCAF8 protein and mRNA expression in several hematologic malignancy and neuroblastoma cell lines (Figure 1A and Supplementary Figure 1A). Two major isoforms of DNMT3A exist: DNMT3A1 (full-length) and DNMT3A2 (short) (Figure 1B). In Kelly (neuroblastoma) cells, both DNMT3A isoforms (DNMT3A1 and 2) and DCAF8 had higher protein expression levels (Figure 1A). We evaluated the protein turnover of DNMT3A1 WT or unstable mutants (W297del and G685R) using ubiquitin-proteasome (UPS) inhibitors such as MG132 (proteasome inhibitor), TAK-243 (MLN7243, ubiquitin E1 enzyme inhibitor), and MLN4924 (neddylation inhibitor). Protein expression was highest in DNMT3A1 WT, followed by G685R, and lowest in W297del [6]. We generated 293T and Kelly cells expressing DNMT3A1 WT or unstable mutants. This plasmid vector produces a DNMT3A1-GFP fusion protein, such that the level of GFP fluorescence detected by flow cytometry allows us to measure the amount of DNMT3A1 protein. Co-expressed DsRed controls for transfection efficiency. We found protein turnover with UPS inhibitors to be inversely proportional to the level of DNMT3A1 expression in the steady state, as UPS inhibitors meaning that the UPS inhibitors had less dramatic effects on WT DNMT3A1 compared to W297del in both 293T and Kelly cells (Figure 1C-D). The stability of DNMT3A WT is due to protection provided by deubiquitinating enzymes and is less susceptible to proteolysis than unstable mutants.

**Figure 1.**
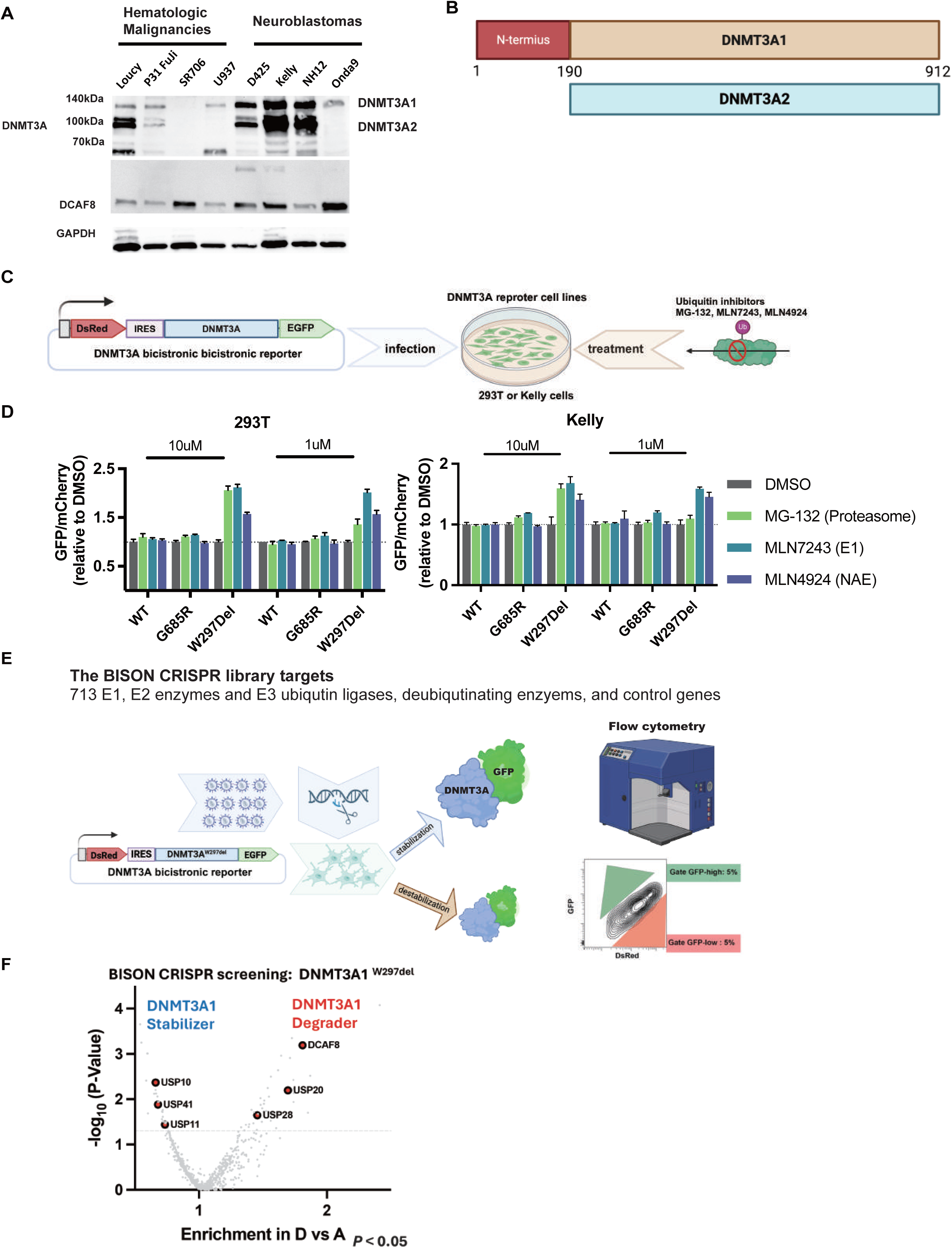
T**a**rgeted **CRISPR screening identifies E3 ligases and deubiquitinating enzymes essential for DNMT3A protein turnover.** A, The image depicts DNMT3A and DCAF8 protein expression followed by western blotting using DNMT3A, DCAF8 and GAPDH antibodies. B, The image depicts the scheme of two major isoforms of DNMT3A exist: DNMT3A1 (full-length) and DNMT3A2 (short) C, The image depicts the scheme of ubiquitin-proteasome inhibitor assay for confirming DNMT3A protein turnover changes. D, The graph depicts the protein expression of DNMT3A WT and unstable mutants (W297del and G685R) with ubiquitin-proteasome inhibitors. Alterations in DNMT3A protein stability after administration of proteasome inhibitor (MG132), a CRL inhibitor (MLN4924), an E1 and ubiquitin enzyme inhibitor (TAK-243) Stability ratio of MFI of DNMT3A-GFP versus MFI of mCherry before and after treatment with the ubiquitin-proteasome inhibitors as measured by flow cytometry 48 hours after transfection. E, Schematic of targeted CRISPR screening to identify ubiquitin modulators essential for DNMT3A protein turnover. Kelly cells were engineered to constitutively overexpress Cas9 and the indicated bicistronic DNMT3AW297Del reporter and then infected with sgRNA libraries targeting ubiquitin ligases. Nine days after infection, we sorted both the top and bottom 5% of cells for DNMT3A-GFP expression. F, the graph depicts the gene enrichment score and P value in targeted CRISPR screening for ubiquitin modulators. DCAF8, USP20, and USP28 genes (red) and USP10, USP11 and USP41 (blue) were enriched and statistically significant.

Since we observed a large change in the expression in USP inhibited DNMT3A1W297Del, we repeated the CRISPR screening to examine factors that mediate DNMT3A1W297Del stability in Kelly cells (Figure 1E). As in our previous study using 293T cells, DCAF8 was enriched as a DNMT3A1W297Del degrader in Kelly cells. We also identified several deubiquitinating enzymes (USP10, USP11 and USP41) as potential stabilizers in the screening (Figure 1F and supplementary Figure 1B-D).

### The ubiquitin specific peptide USP11 induced DNMT3A deubiquitination and stabilization

Protein-protein interactions play a crucial role in protein turnover [14]. E3 ligases bind to target proteins and induce their ubiquitination and degradation. In contrast, deubiquitinating enzymes bind to substrate proteins and prevent ubiquitination and degradation. To better understand DNMT3A protein turnover and which deubiquitinating enzymes interact with DNMT3A, we used IP-MS targeting DNMT3A1W297del. We analyzed the interactome between DNMT3A1W297del and deubiquitinating enzymes with or without the E1 ubiquitin inhibitor TAK-243 (MLN7243), which stabilizes DNMT3A1W297del (Figure 2A). Several deubiquitinating enzymes (USP1 and USP11) increased their interaction with DNMT3A1W297del after deubiquitylation (Figure 2B and supplementary Figure 2A-C). Both CRISPR and IP-MS screenings identified USP11 as a stabilizer of mutant DNMT3A, indicating that it is a strong DNMT3A deubiquitinating enzyme (Figure 1E and Figure 2 B).

**Figure 2.**
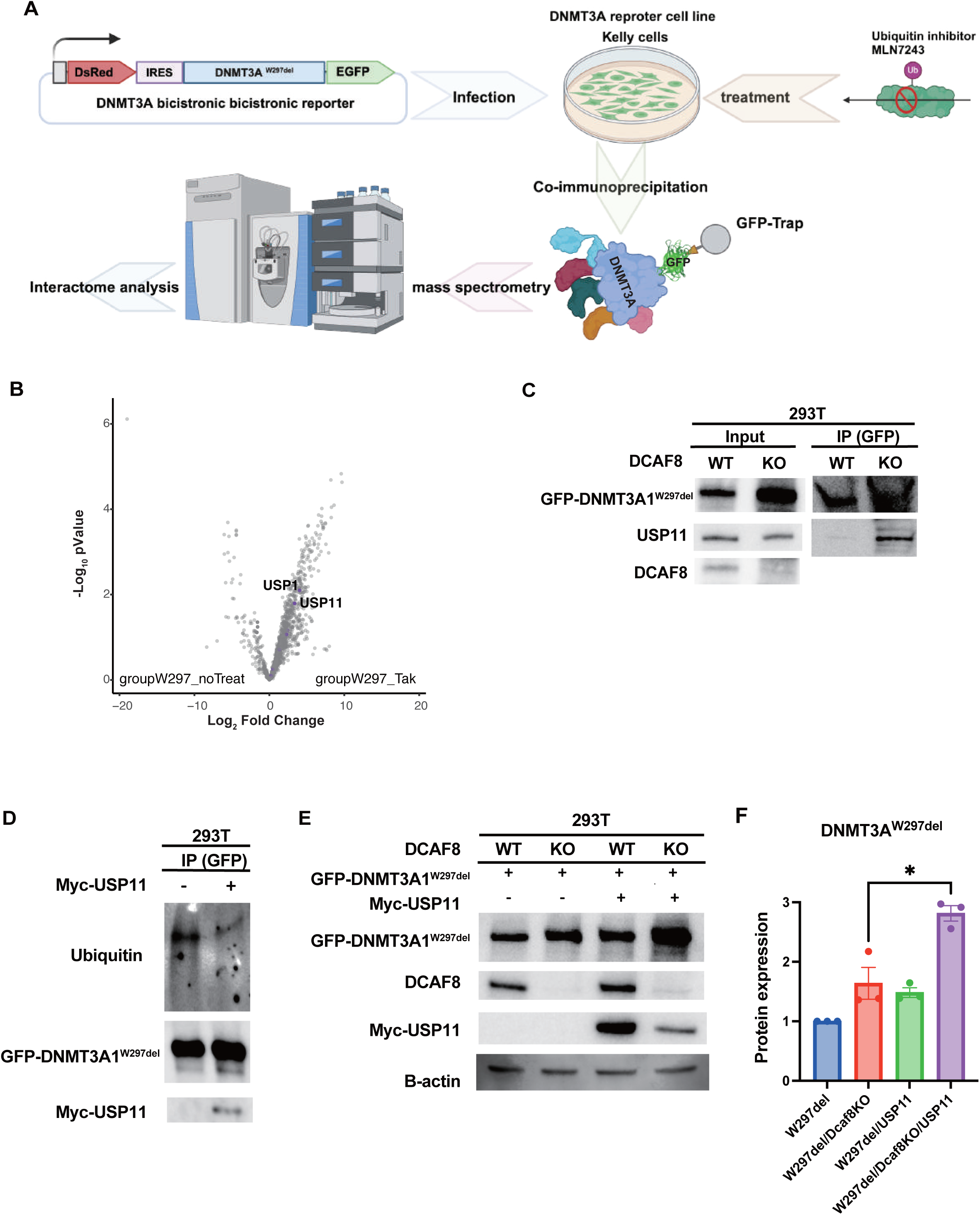
U**S**P11 **regulated DNMT3A protein turnover countering protein degradation by FCAF8** A, Schematic of immunoprecipitation (IP) mass spectrometry to identify deubiquitinating enzymes essential to interact with DNMT3A. Kelly cells were engineered to constitutively overexpress the indicated bicistronic GFP-DNMT3AW297Del reporter and treated with TAK-243. To subjects cell lysate was performed I P and mass spectrometry analysis. B, The graph depicts the enriched score of the interaction between DNMT3AW297del and deubiquitinating enzymes with or without 1nM TAK-243 for 6 hours. C. 293T cells were transfected with GFP-DNMT3A1W297Del. Cell lysates were immunoprecipitated with anti-GFP antibody, following the detection of USP11-bound DNMT3A1W297Del with anti-USP11 antibody. D. 293T cells were transfected with GFP-DNMT3A1W297Del. Cell lysates were immunoprecipitated with anti-GFP antibody, following the detection of DNMT3A1W297Del ubiquitination with anti-ubiquitin antibody. E, F. In the presence or absence of DCAF8, 293T cells were transfected with GFP-DNMT3A1W297Del and/or Myc-USP11. Cell lysates were immunoblotted following the detection of DNMT3A1W297Del, USP11 and DCAF8 protein expression with anti-GFP-Myc and-DCAF8 antibodies. F: The graph depicts the protein expression in Figure 2E (data are shown as the mean ± S.E.M, *, p = 0.0310) The W297del samples were set as 1.

We next examined whether DCAF8 was involved in the interaction between DNMT3A1W297Del and USP11. We introduced GFP-DNMT3A1W297Del into 293T cells in the presence or absence of DCAF8. Cell lysates were subjected to IP with GFP trap [15], followed by immunoblotting with anti-USP11 to detect DNMT3A1W297Del interacting with USP11. We found that USP11 strongly interacted with GFP-DNMT3A1W297Del in DCAF8 KO 293T cells (Figure 2C). Next, we introduced GFP-DNMT3A1W297Del and Myc-USP11 into 293T cells to determine whether USP11 induces DNMT3A1W297Del deubiquitination. Cell lysates were subjected to IP with GFP trap, followed by immunoblotting with anti-ubiquitin to detect ubiquitinated DNMT3A1W297Del. We found that USP11 transduction into 293T cells decreased ubiquitinated DNMT3A1W297Del (Figure 2D). Finally, we investigated how DCAF8 influences the effect of USP11 on DNMT3A1W297Del protein expression by overexpression of USP11 in DCAF8 KO 293T cells.

We found that USP11 overexpression increased DNMT3A1W297del protein expression more in DCAF8 KO 293T cells than DCAF8 WT cells (Figure 2E-F). Taken together, these results suggest that USP11 is a major deubiquitinating enzyme for DNMT3A protein turnover by countering ubiquitination and degradation by DCAF8.

### Inhibiting DNMT3A ubiquitination restores the mislocalization of DNMT3A1W297del

DNMT3A has three functional domains: Pro-Trp-Trp-Pro (PWWP), ATRX-DNMT3A-DNMT3L (ADD), and MTase [16]. Previous studies showed that several PWWP mutations, including W297del, are associated with DNMT3A protein instability and dysregulated localization from nucleus to cytoplasm [17]. Thus, we investigated whether DNMT3A1W297del deubiquitination could restore the localization of DNMT3A1W297del to the nucleus. We generated 293T cells expressing DNMT3A1 WT or W297del and performed immunofluorescence analysis to assess localization (Figure 3A-C and Supplementary Figure 3A-B). We extracted cytoplasmic, membrane, and nuclear proteins from transduced cells and found that DNMT3A1W297del expression decreased in nucleus and increased cytoplasm and membrane (Figure 3D). Similar to the previous report [17], W297del mutant sequestered DNMT3A1 protein toward the cytoplasm, resulting in a more diffuse localization pattern.

**Figure 3.**
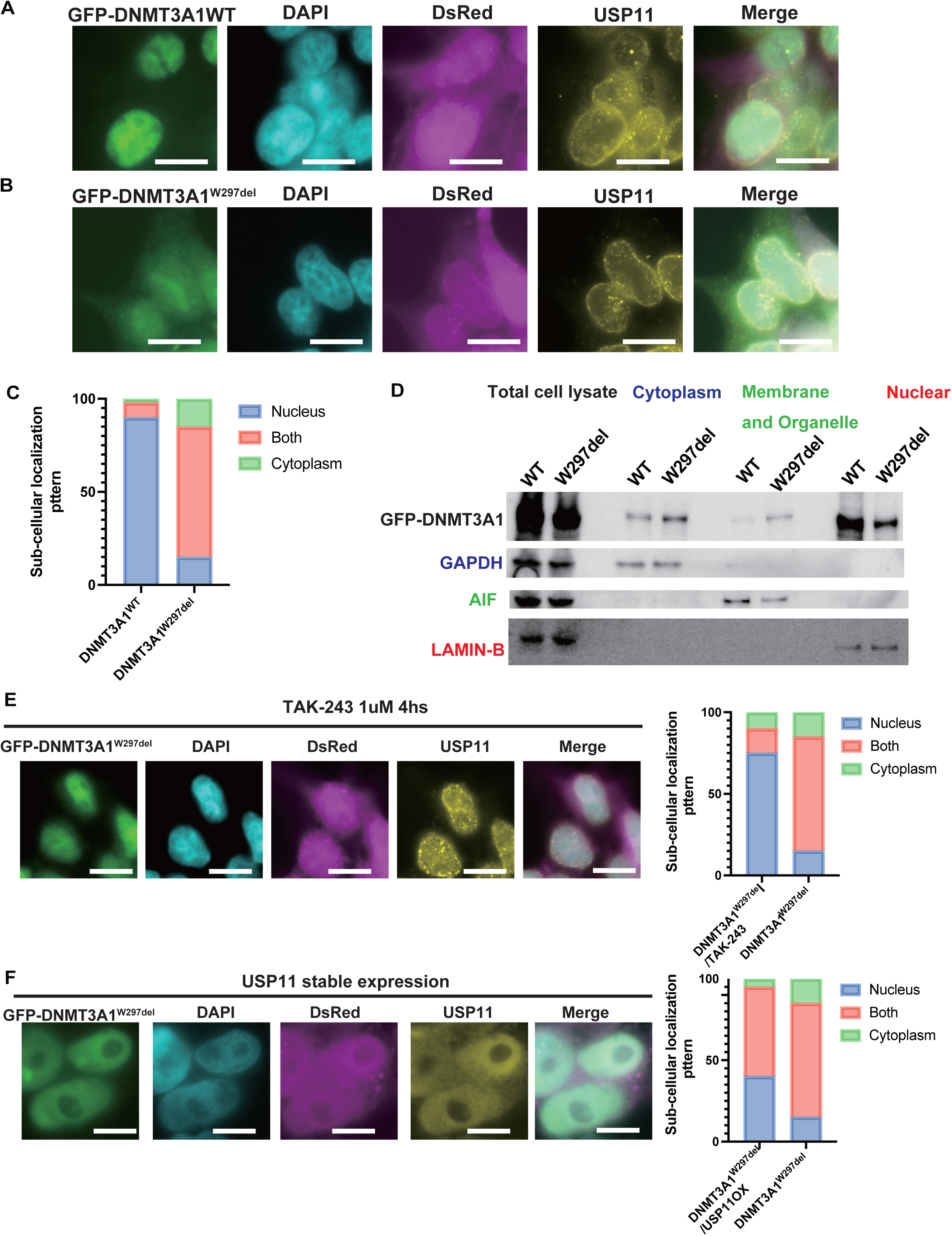
D**N**MTA **deubiquitination restored the mislocalization of DNMT3AW297del** A, B 293T cells were transfected with GFP-DNMT3A WT or-W297del and were stained anti-USP11 (rabbit) antibodies followed by anti-rabbit Alexa 647 (yellow). DNMT3A expression detected GFP (Green), nuclei and cytoplasm detached, DsRed (Red). Nuclei were visualized with DAPI (Blue). Laser scanning microscopy (Keyence) was used for the imaging. C, The graph depicts the subcellular localization pattern of GFP-DNMT3A WT or-W297del. D, 293T cells were transfected with GFP-DNMT3A WT or-W297del. nuclear, membrane/organelle and cytoplasmic fractions were isolated, following the detection of DNMT3A with anti-GFP antibody. GAPDH was used as a loading control for cytoplasm, AIF for membrane/organelle, and LAMIN-B for nuclear proteins. E, 293T cells were transfected with GFP-DNMT3A1W297del and were stained anti-USP11 (rabbit) antibodies followed by anti-rabbit Alexa 647 (yellow) with or without 1nM of TAK-243 for 6 hours. DNMT3A expression detected GFP (Green), nuclei and cytoplasm detached, DsRed (Red). Nuclei were visualized with DAPI (Blue). Laser scanning microscopy (Keyence) was used for the imaging. The graph depicts the subcellular localization pattern of GFP-W297del with or without TAK-243. F, 293T cells were transfected with GFP-W297del and USP11 were stained anti-USP11 (rabbit) antibodies followed by anti-rabbit Alexa 647 (yellow), DNMT3A expression detected GFP (Green), nuclei and cytoplasm detected DsRed (Red), nuclei were visualized with DAPI (Blue). Laser scanning microscopy (Keyence) was used for the imaging. The graph depicts the subcellular localization pattern of GFP-W297del with or without USP11 overexpression.

To understand if E1 ubiquitin enzymes play a role in the localization of DNMT3A1 W297del, we cultured DNMT3A1W297del 293T cells with TAK-243, an E1 ubiquitin enzyme inhibitor. We found that DNMT3A1 W297del localization was restored from cytoplasm to nucleus in the presence of TAK-243 (Figure 3E, Supplementary Figure 3C). Finally, we evaluated whether USP11 was involved in DNMT3A1W297del subcellular localization. We generated 293T cells expressing W297del with USP11 transduction and performed immunofluorescence analysis (Figure 3F and Supplementary Figure 3D). We found that USP11 also impacts DNMT3A subcellular localization by transducing USP11 into DNMT3A1W297del 293T cells. This study revealed that endogenous USP11 is predominantly localized to the nucleus and nuclear membrane, while USP11 overexpression is localized to the nucleus and cytoplasm. Finally, we confirmed USP11 overexpression modesty restored DNMT3AW297del localization to the nucleus, possibly due to the subcellular localization of USP11 (Figure 3F and Supplementary Figure 3D). Thus, the ubiquitin system regulates DNMT3A1 mislocalization, which is induced the PWWP domain mutations and blocking DNMT3A1W297del ubiquitination can partially restore DNMT3A1W297del subcellular localization from the cytoplasm to the nucleus.

### USP11 stimulates DNMT3A1 SUMOylation and facilitates its protein turnover

To understand the regulation of DNMT3A protein turnover, we examined the relationship between DNMT3A WT and USP11. First, we performed co-IP-MS to compare which proteins differentially interact with DNMT3A1 WT or W297del (Figure 4A and Supplementary Figure 4A-C). We found that proteins involved in SUMOylation strongly and preferentially interacted with DNMT3A1 WT (Figure 4B and Supplementary Figure 4D). SUMOylation is a post-translational modification that plays a role in protein stability, nuclear-cytosolic transport, and transcriptional regulation [18]. We also performed co-IP-MS with W297del in the presence or absence of DCAF8 and saw that W297del preferentially interacted with SUMO proteins in DCAF8 KO cells (Figure 4C and Supplementary Figure 4E). Overall, this study suggests that unstable DNMT3A loses its interaction with SUMO proteins, which can be rescued by DCAF8 KO.

**Figure 4.**
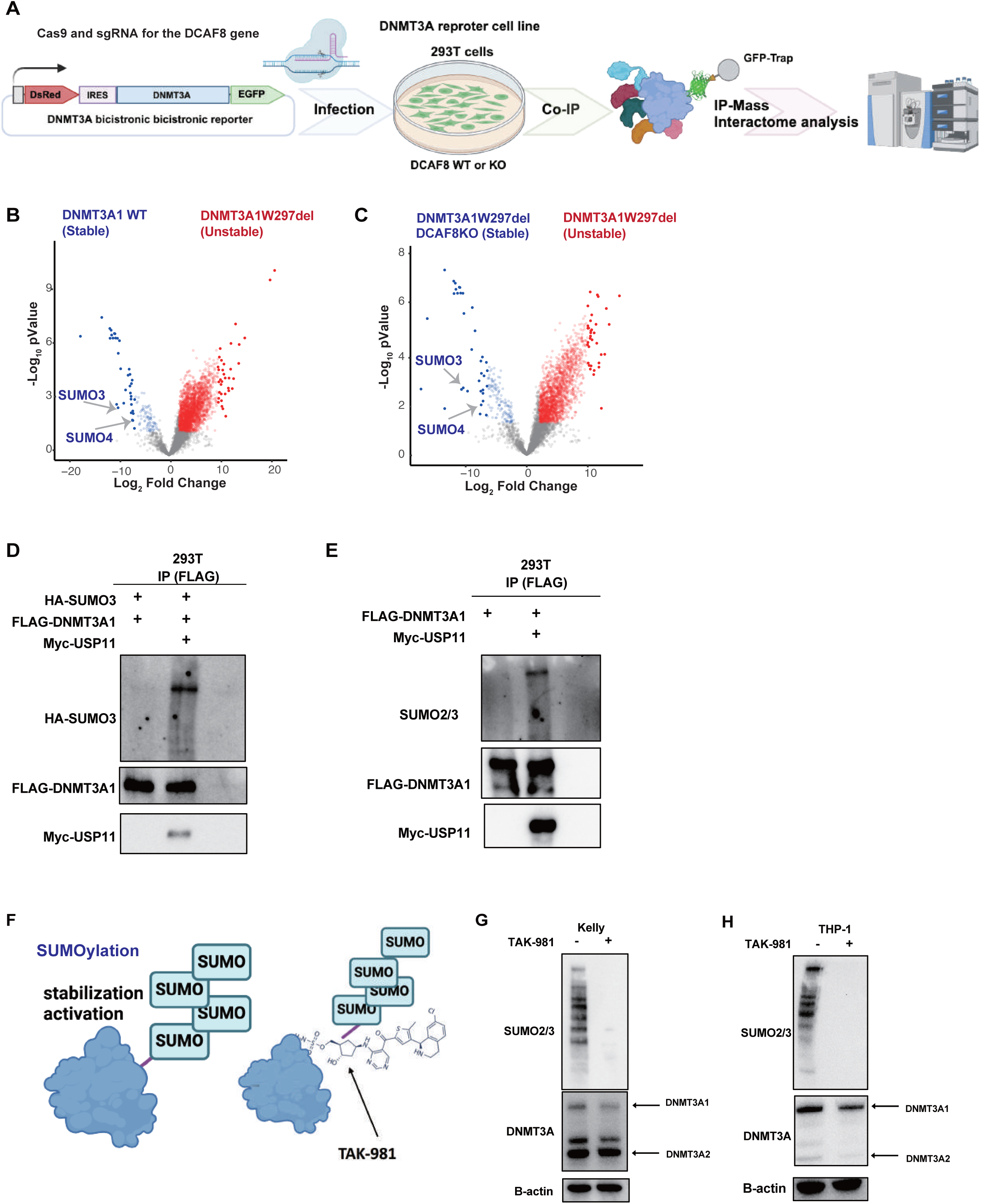
U**S**P11 **induced DNMT3A SUMOylation and its essential for DNMT3A protein stability** A. Schematic of immunoprecipitation (IP) mass spectrometry to identify comparing DNMT3A stable (DNMT3A WT or W297Del in the absence of DCAF8) or an unstable protein (W297Del) interactome. 293T cells were engineered to constitutively overexpress the indicated bicistronic GFP-DNMT3A WT or W297Del reporter in the presence or absence of DCAF8. To subjects cell lysate was performed I P and mass spectrometry analysis. B, C. The graph depicts the enriched score of the interaction proteins with stable DNMT3A or unstable DNMT3A. D. 293T cells were transfected with FLAG-DNMT3A1, Myc-USP11 and HA-SUMO3. Cell lysates were immunoprecipitated with anti-FLAG (DNMT3A) antibody, following the detection of DNMT3A1 SUMOylation with anti-HA antibody. E. 293T cells were transfected with FLAG-DNMT3A1, Myc-USP11. Cell lysates were immunoprecipitated with anti-FLAG (DNMT3A) antibody, following the detection of DNMT3A1 endogenous SUMOylation with anti-SUMO2/3 antibody. F. The scheme depicts TAK-981, a first-in-class Inhibitor of SUMO-Activating enzymes. G, H. Kelly or THP-cells were cultured with 1uM ofTAK-981 for 24 hours. Cell lysates were detected with anti-DNMT3A,-SUMO2/3 and-B-actin antibodies.

To test the hypothesis that SUMOylation is important to DNMT3A protein stability and turnover, we examined whether USP11 is involved in DNMT3A1 SUMOylation. We transduced FLAG-DNMT3A1, Myc-USP11 and HA-SUMO3 into 293T cells. Cell lysates were subjected to IP with FLAG (DNMT3A1) antibody, followed by immunoblotting with anti-HA (SUMO3) to detect DNMT3A SUMOylation (Figure 4D). Additionally, we evaluated endogenous DNMT3A SUMOylation by subjecting cell lysates from FLAG-DNMT3A1 and Myc-USP11-transduced 293T cells to immunoprecipitation with FLAG (DNMT3A1), followed by immunoblotting with anti-SUMO3 (Figure 4E). We confirmed that USP11 induced DNMT3A1 SUMOylation, suggesting that DNMT3A1 protein stability and SUMOylation are maintained by UPS11 (Figure 4D, E). We next examined how DNMT3A SUMOylation impacts its protein turnover. We cultured Kelly and THP-1 cell lines with TAK-981, a first-in-class Inhibitor of SUMO-Activating Enzyme [19](Figure 4F). We found that TAK-981 strongly reduced DNMT3A1 but not DNMT3A2 protein levels (Figure 4G, H). These studies suggest that DNMT3A is stabilized by interaction with SUMO proteins. Protein SUMOylation is essential for SUMO E3 ligase interaction with target proteins [18, 20]. Thus, we examined whether USP11 enhances the DNMT3A1-SUMO E3 ligase interaction. We transduced GFP-DNMT3A1, and Myc-USP11 into 293T cells and subjected cell lysates to immunoprecipitation with GFP trap before performing MS analysis (Supplementary Figure 5A). This study showed that USP11 promotes the binding of multiple E3 SUMO ligases (CBX4 and PIAS1) to DNMT3A1 (Supplementary Figure 5B). Target protein residues previously identified as SUMOylated contain the consensus motif KxE [21]; x is any amino acid. Interestingly, we found the motif to be abundant at the N-terminus of DNMT3A, a feature unique to DNMT3A1. (Supplementary Figure 5C). Taken together, USP11 promotes the deubiquitination and SUMOylation of DNMT3A1 and SUMO E3 ligases may accelerate DNMT3A1 SUMOylation and protein turnover.

### USP11 enhances the binding of DNMT3A1 to the polycomb complex as well as the DNA Mtase activity of DNMT3A1

We further examined USP11’s relationship with DNMT3A1 and its effects on DNMT3A1 function. We first performed co-IP to compare USP11’s interaction with DNMT3A1 and DNMT3A2. We transduced Myc-USP11 and FLAG-DNMT3A1 or FLAG-DNMT3A2 into 293T cells and subjected lysates to IP with FLAG (targeting DNMT3A1 or 2) antibody, followed by immunoblotting with anti-Myc (USP11). We found that USP11 strongly interacted with DNMT3A1 and the N-terminus of DNMT3A might be necessary for its interaction with USP11, because of DNMT3A2 luck of the DNMT3A N-terminus (Figure 5A, Supplementary Figure 6A, B). To determine whether endogenous USP11 controls DNMT3A protein expression in Kelly cells, we transduced Cas9 protein and USP11-targeting sgRNA into Kelly cells and performed WB for DNMT3A. Depletion of USP11 decreased DNMT3A1 protein expression (Supplementary Figure 6C-E).

**Figure 5.**
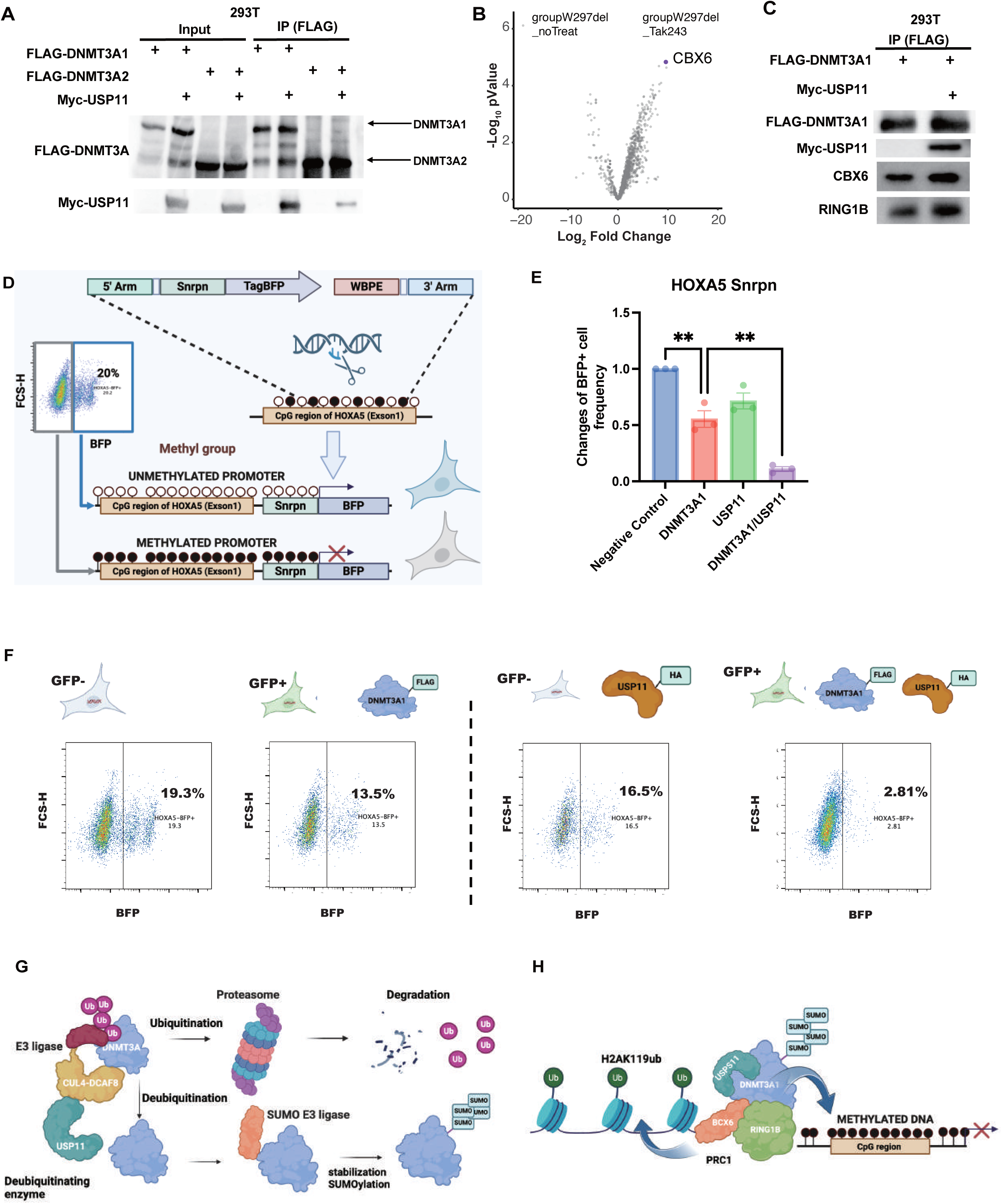
U**S**P11 **induced DNMT3A SUMOylation and its essential for DNMT3A protein stability** A, 293T cells were transfected with FLAG-DNMT3A1 or 2 and Myc-USP11. Cell lysates were immunoprecipitated with anti-FLAG (DNMT3A) antibody, following the detection of DNMT3A interacted USP11 with anti-myc antibody for western blotting. B, The graph depicts the enriched score of the interaction between DNMT3AW297del and deubiquitinating enzymes with or without 1nM TAK-243 for 6 hours. C, 293T cells were transfected with FLAG-DNMT3A1 and Myc-USP11. Cell lysates were immunoprecipitated with anti-FLAG (DNMT3A) antibody, following the detection of DNMT3A interacted USP11 with anti-myc, CBX6 with anti-CBX6, RING1B with anti-RING1B antibodies for western blotting. D, DNMT3A DNA Methyltransferase activity analysis using the HOXA5-Snrpn-BFP methylation. Knocking in (KI) Snrpn promoters to the HOXA5 locus in reverse orientation in 293T cells linked to BFP and bovine polyA signals. E, F, Methyltransferase activity assay. The figure depicts blue fluorescence intensity in the FLAG-DNMT3A and/or HA-USP11 transduced cells as measured by flow cytometry on day 10. The graphs depicted the BFP positive cells. The negative control frequency of BFP+ cells was set as 1. One-way ANOVA with Tukey’s multiple comparisons was completed. *p < 0.05, **p < 0.01 *** p < 0.001 ****p < 0.0001.

Recent studies suggest that DNMT3A1 regulates polycomb complex functions by interacting with polycomb complexes and Histone H2A [13, 22]. To determine whether DNMT3A1W297del interacts with members of the polycomb complex, we compared the interactomes of DNMT3A1W297del with or without TAK-243, which we found to stabilize DNMT3A1W297del (Figure2A, 5B and Supplementary Figure 2C). Indeed, we found that W297del in the presence of TAK-243 strongly interacted with CBX6 (Figure 5B), a component of the polycomb complex. We also examined whether USP11 enhances the interaction between DNMT3A1 and polycomb complex proteins by introducing GFP-DNMT3A1W297Del and Myc-USP11 into 293T cells and performing IP with GFP trap on lysates from these cells. We found that USP11 enhanced the interaction of DNMT3A1 with polycomb complex proteins CBX6 and RING1B (Figure 5C). Overall, USP11 strongly interacted with DNMT3A1 but not with DNMT3A2 (Figure 5A), and USP11 was important in the binding of DNMT3A1 to the polycomb complex.

Finally, we examined whether USP11 is involved in the Mtase activity of DNMT3A1 using the HOXA5-Snrpn-blue fluorescence (BFP) reporter system [23, 24]. We inserted a Snrpn-BFP reporter into the HOXA5 promoter locus in 293T cells such that increased DNA methylation at HOXA5-Snrpn-BFP would decrease BFP levels, detectable by flow cytometry (Figure 5D). To test the fidelity of this assay, we transduced the cells with a retrovirus vector carrying HA-USP11 co-expressing a puromycin resistance gene. Puromycin was used to create stable USP11 expression strains, which were then transduced with a retrovirus vector carrying DNMT3A1 co-expressing GFP. Following the introduction of DNMT3A1, the percentage of BFP-positive cells significantly declined (Figure 5E, F). USP11 alone reduced BFP-positive cells slightly less than DNMT3A alone. On the other hand, co-expression of DNMT3A1 and USP11 dramatically reduced BFP expression (Figure 5E, F and Supplementary Figure 7A-C). Based on these findings, USP11 strongly increases the DNA Mtase activity of DNMT3A1 in the HOXA5 prompter region.

Overall, we found that USP11 induces DNMT3A deubiquitylation and stabilization, which counteracts degradation by DCAF8. In addition, USP11 accelerates DNMT3A SUMOylation and this SUMOylation is essential for maintaining DNMT3A protein stability (Figure 5G). Further, USP11 enhances the binding of DNMT3A1 to the polycomb complex and ensures DNMT3A DNA Mtase activity (Figure 5H).

## Discussion

Here we characterize the mechanisms of DNMT3A protein turnover — a new biological pathway with therapeutic potential for CH and TBRS. Using CRISPR screening and IP-MS, we identified several deubiquitinating enzymes, including USP11, that stabilize DNMT3A. Our CRISPR screen identified DCAF8 as a degrader of DNMT3A, consistent with our previous findings [6], and we found that USP11 counters DCAF8 to stabilize DNMT3A. Therefore, USP11 and DCAF8 play opposite roles in DNMT3A protein turnover. A previous report indicated that DCAF8 and USP11 regulate myeloid leukemia factor (MLF) expression in opposite directions, which mean degradation of MLF leads to DCAF8 and stabilization by USP11, mirroring our results in DNMT3A turnover [25]. However, why USP11 and DACF8 are specifically involved in the regulation of proteolysis remains unclear. We identified other E3 ligases and deubiquitinating enzymes such as RNF26, USP1, USP10 and USP41, which may also be involved in the protein turnover of DNMT3A, requiring further investigation.

We found that USP11 induced DNMT3A SUMOylation and enhanced the interaction between DNMT3A and E3 SUMO ligases such as CBX4 and PIAS1. Previous reports indicated that SUMOylation increased DNMT1 DNA MTase activity [26], and SUMOylation also enhances DNMT family DNA binding efficiency [27]. Protein-DNA binding is important to protein stabilization [28], suggesting that USP11 may contribute to protein stabilization and deubiquitination by accelerating DNA binding of DNMT3A through SUMOylation. Previous research suggests that PIAS1 knockout mice exhibit abnormal differentiation of HSCs caused by Dnmt3a DNA MTase deficiency [29], indicating that DNMT3A SUMOylation is essential for its role in controlling HSC differentiation. We used a cell-based assay to confirm that USP11 increases DNMT3A DNA MTase activity, and validating this finding in animal models is an important future direction. We also demonstrated that USP11 is involved in DNMT3A1 binding to the polycomb complex. Recent studies have shown that DNMT3A1 may regulate H2AK119ub as well as the polycomb complex in the bivalent domains of chromatin [13, 22, 30]. In addition, patients with AML who have DNMT3A unstable mutations show abnormal DNA methylation in the bivalent domains [10]. However, it remains unclear how DNMT3A protein interacts with the chromatin complex including the modification of H2AK119ub via the polycomb complex and DNA methylation in the bivalent domains. A recent study showed that DNMT3A is normally recruited to intergenic regions by H3K36me2, but DNMT3AW297Del unstable mutants fail to bind to chromatin [31]. Nuclear receptor binding SET domain protein1 (NSD)-mediated H3K36me2 is required for the recruitment of DNMT3A and maintenance of DNA methylation [32]. Interestingly, TBRS shares clinical features with Sotos syndrome which is caused by haploinsufficiency of NSD1 [31]. NSD1 and H3K36me2 may be related to the regulation of the chromatin complex and DNMT3A via its protein stability. Thus, it is necessary to investigate the relationship between DNMT3A protein turnover and histone modification further.

In many CH-related DNMT3A mutations, as well as some overlapping TBRS mutations, DNMT3A protein expression is downregulated. Therefore, unstable DNMT3A mutations may lead to the development of MDS, AML, and TBRS and its protein turnover regulation may be a worthwhile therapeutic target. We showed that TAK-243, an E1 ubiquitin ligase inhibitor, and USP11 could restore DNMT3A protein expression and proper nuclear localization. These data suggest that DNMT3A deubiquitination is one potential strategy to modulate DNMT3A protein turnover. TAK-243 has shown therapeutic effects in several human cancer xenograft mouse models [33]. Future research should examine the therapeutic and off-target effects of TAK-243 on unstable DNMT3A in vivo. Another approach might be inhibition of DCAF8, so the development of DCAF8 inhibitors may be useful. As TAK-243 and DCAF8 have multiple targets, evaluating potential side effects is paramount and development of DNMT3A-targeted deubiquitinase-targeting chimeras (DUBTACs) may be necessary.

DUBTACs directly induce protein interaction between a target protein and a deubiquitinating enzyme [34]. As a result, the target protein is subject to deubiquitination and the protein is stabilized. Proteolysis Targeting Chimeras (PROTACs), on the other hand, induce interaction between a target protein and an E3 ligase, resulting in ubiquitination and degradation [35]. Because this approach degrades the target protein directly, there are fewer side effects. Another advantage of this strategy is that PROTACs and DUBTACs can regulate protein turnover of target proteins, which is not limited to endogenous target protein degradation mechanisms. As unstable DNMT3A mutants other than DNMT3AW297del are likely regulated for degradation by proteins other than USP11 and DCAF8, DNMT3A-targeted DUBTACs may be effective in treating 30% of DNMT3A-CH [6]. USP11 also increases DNMT3A DNA MTase activity, so creating DNMT3A-USP11 specific DUBTAC could be an effective treatment for CH and other diseases such as MDS, AML and TBRS.

In this study, we reveal a new molecular mechanism for DNMT3A protein turnover through USP11 and SUMOylation. By modulating DNMT3A function, USP11 may play a significant role in HSC regulation. Additionally, USP11 is strongly associated with DNMT3A1, which plays an important role in neural development by regulating neural gene expression. Therefore, USP11 may play an important role in neural development. As a result, activating USP11 may be useful in the prevention and treatment of diseases such as CH and TBRS caused by unstable DNMT3A mutations.

## Methods Cell culture

Kelly, THP-1, Loucy, P31, Fuji SR706 and U937 cells were cultured in RPMI-1640 with 10% fetal bovine serum (FBS) and 1% penicillin-streptomycin. 293T, D425, NH12 and Onda9 cells were cultured in DMEM with 10% fetal bovine serum (FBS) and 1% penicillin-streptomycin.

### Plasmids

For DNMT3A expression, FLAG-tagged DNMT3A1 or 2 in a pCAG vector and GFP-tagged DNMT3A1 in a pLenti vector were used. For Myc-tagged USP11 and HA-tagged SUMO3 in a pCDNA3 vector were used for their expression. FLAG-tagged DNMT3A1 in a pMYs-IRES-GFP (pMYs-IG) vector were used to express DNMT3A1, and HA-tagged USP11 in a pMYs-IRES-Puro vector were used to express USP11.

### CRISPR-DNMT3A Stability Screen

The human BISON CRISPR-KO library, which contains 2,852 guide RNAs, targets 713 E1, E2, E3, deubiquitinases, and control genes [11]. It was cloned into the pXPR003 as previously described [36] by the genome perturbation platform (Broad Institute). The lentivirus particles for the library were produced in a T-175 flask format. Briefly, 18 °ø 106 Kelly cells were seeded in 25 mL DMEM supplemented with 10% FBS and penicillin/streptomycin/glutamine. The next day, a packaging mix was prepared: 40 μg psPAX2, 4 μg pVSV-G, and 32 μg of the library in 1 mL OptiMem (Invitrogen) and incubated for 5 minutes at room temperature. This mix was combined with 244 μL TransIT-LT1 (Mirus) in 5 mL OptiMem, incubated for 30 minutes at room temperature, and then applied to cells. Two days post transfection, cell debris was removed by centrifugation. The lentivirus particles containing medium were collected and stored at −80°C before use. Kelly cells were engineered with constitutively expressing Cas9 and bicistronic DNMT3AW297Del reporter. Then, 2 °ø 106 engineered 293T cells were added with 10% (v/v) of the human BISON CRISPR-KO in 2 mL medium and spin-infected (2,400 rpm, 2 hours, 37°ΔC). Twenty-four hours post infection, sgRNA-infected cells were selected with 2 μg/mL puromycin for 2 days. On the ninth day postinfection, populations were separated using fluorescence activated cell sorting. Two populations were collected (top 5% and lowest 5%) based on the eGFPDNMT3A to mCherry MFI ratio. Sorted cells were harvested by centrifugation and subjected to direct lysis buffer reactions (1 mmol/L CaCl2, 3 mmol/L MgCl2, 1 mmol/L EDTA, 1% Triton X-100, Tris pH 7.5, with freshly supplemented 0.2 mg/mL proteinase). The sgRNA sequence was amplified in a first PCR reaction with eight staggered forward primers. Then, 20 μL of direct lysed cells was mixed with 0.04 U Titanium Taq (Takara Bio 639210), 0.5 °ø Titanium Taq buffer, 800 μmol/L dNTP mix, 200 nmol/L P5-SBS3 forward primer, and 200 nmol/L SBS12-pXPR003 reverse primer in a 50-μL reaction. The samples were heat activated at 94°ΔC for 5 minutes; kept at 94°ΔC for 30 seconds, 58°ΔC for 10 seconds, and 72°ΔC for 30 seconds and repeated from the second step for 15 cycles; and heated at 72°ΔC for 2 minutes. Then, 2 μL of the primary PCR product was used as the template for 15 cycles of the secondary PCR, where Illumina adapters and barcodes were added [0.04 U Titanium Taq (Takara Bio 639210), 1 °ø Titanium Taq buffer, 800 μmol/L dNTP mix, 200 nmol/L SBS3-Stagger-pXPR003 forward primer, 200 nmol/L P7-barcode-SBS12 reverse primer]. An equal amount of all samples was pooled and subjected to preparative agarose electrophoresis followed by gel purification (Qiagen). Eluted DNA was further purified by NaOAc and isopropanol precipitation. Amplified sgRNAs were quantified using the Illumina NextSeq platform. Read counts for all guides targeting the same gene were used to generate P values. The data analysis pipeline comprised the following steps: (i) Each sample was normalized to the total read number. (ii) For each guide, the ratio of reads in the stable versus unstable sorted gate was calculated, and guide RNAs were ranked. (iii) The ranks for each guide were summed for all replicates. (iv) The gene rank was determined as the median rank of the four guides targeting it. (v) P values were calculated, simulating a corresponding distribution over 100 iterations.

### Transfection, western blotting, and immunoprecipitation

293T cells were transiently transfected with 5 ug of vector, GFP or FLAG-tagged DNMT3A, MYC-tagged USP11, or HA-tagged SUMO3 mixed with lipofectamine 2000. The cells were cultured for 48 h after the transfection and lysed in Hypotonic buffer and Lysis buffer (see below). For FLAG-targeted immunoprecipitation, the cell lysates were incubated with anti-FLAG M2 (SIGMA, Louis, MO, USA, Catalog Number F3165) antibody for 15 min at 25°C. Then the samples were incubated with Dynabeads protein-G (Thermo Fisher Scientific, Waltham, MA, USA) for 15 min at 25°C. For GFP-targeted immunoprecipitation the cell lysates were incubated GFP trap (Protein Tek GFP-Trap® Magnetic Agarose, IL, USA, Catalog Number gtma-20) for 60 min at 4°C The precipitates were washed three times with cell lysis buffer (Cell Signaling Technology, Danvers, MA, USA, Catalog Number 9803) containing 1 mM phenylmethanesulfonylfluoride fluoride, subjected to sodium dodecyl sulfate-polyacrylamide gel electrophoresis (SDS-PAGE), and analyzed by western blotting. For some experiments, we isolated nuclear and cytoplasmic fractions before the immunoprecipitation (see below). Western blotting was performed with anti-FLAG M2-Peroxidase (Sigma, Catalog Number A8592), anti-Myc 9E10 (Santa Cruz Biotechnology, sc-40), anti-HA-Peroxidase (Roche, 12CA5), anti-GFP (NOVUS, Catalog Number NB600-308), anti-USP11 (Abcom, Catalog Number ab109232), anti-Lamin B1 (Santa Cruz Biotechnology, Catalog Number B1304), anti-GAPDH (Cell Signaling Technology, Catalog Number 5174), anti-AIF (Cell Signaling Technology, Catalog Number 5318), Anti-DCAF8 Bethyl lab, Catalog Number A301-556A), anti-B-actin (Cell Signaling Technology, Catalog Number, 3700S), and anti-SUMO2/3 (Cell Signaling Technology, Catalog Number, 4971). Signals were detected by using Clarity Western ECL Substrate (BioRad, Catalog Number, 170-5060), and immunoreactive bands were visualized using a ChemiDoc MP imaging System (BioRad,Catalog Number 12003154). Band intensities were measured using ImageLab (BioRad).

### Immunofluorescence analysis

293T cells introduced with control vector, GFP-tagged DNMT3A, which is coexpressed DsRed were fixed with 2% paraformaldehyde, permeabilized with 0.2% Triton X-100, blocked with 2% BSA, and then incubated with anti-GFP (NOVUS, Catalog Number NB600-308) and anti-USP11 (Abcom, Catalog Number ab109232) antibodies, followed by labeling with Alexa Fluor 568– conjugated anti-rabbit antibody (Thermo Fisher Scientific, Catalog Number A11011) and Alexa Fluor 488–conjugated anti-mouse antibody (Thermo Fisher Scientific, Catalog Number A11034). Nuclei were visualized with DAPI (Bio Legend). Fluorescent images were analyzed on a Keyence BZ-X800 microscope (Keyence).

### Viral transduction

Retroviruses for human cells and lentiviruses were produced by transient transfection of 293T cells with viral plasmids along with gag-, pol-, and env-expressing plasmids using the calcium-phosphate method [37]. Retrovirus transduction to the cells was performed using Retronectin (Takara Bio Inc., Otsu, Shiga, Japan).

### IP Mass spectrometry

The cells were lysed in 3 volumes of NETN buffer (50mM Tris pH 7.3, 170mM NaCl, 1mM EDTA, 0.5% NP-40) using probe sonication followed by ultracentrifugation at 44,000rpm for 20min at 4oC. The lysate was incubated with 20µl of GFP-Trap agarose beads (Chromotek gta) for 1hr at 4oC. The beads were briefly washed with lysis buffer, boiled in 2X NUPAGE® LDS sample buffer (Invitrogen) and subjected to SDS-PAGE (NuPAGE 10% Bis-Tris Gel, Invitrogen). In-gel protein digestion of the whole lane was done using trypsin and LysC protease mix. The LC-MS/MS analysis was carried out using a nanoLC1000 system coupled to Orbitrap Fusion mass spectrometer (Thermo Scientific, San Jose, CA). The MS raw data was searched in Proteome Discoverer 2.1 software (Thermo Scientific, San Jose, CA) with Mascot algorithm (v2.4, Matrix Science) against the human NCBI refseq database (updated 2021_12_23). The peptides identified from the mascot result file were validated with 5% false discover rate (FDR). The gene product inference and quantification were done with label-free iBAQ approach using the ‘gpGrouper’ algorithm [PMID: 30093420]. The differentially expressed proteins were calculated using the moderated t-test to calculate p-values and log2 fold changes in the R package limma [PMID: 25605792]. For statistical assessment, missing value imputation was employed through sampling a normal distribution N(μ-1.8 σ, 0.8σ), where μ, σ are the mean and standard deviation of the quantified values.

### HOXA5-Snrpn-BFP Methylation Reporter Analysis

HOXA5-Snrpn-BFP Methylation Reporter Analysis DNA for the Snrpn promoter, BFP (tagBFP), woodchuck hepatitis virus post translational regulatory elements, bovine growth hormone polyadenylation signal, and HOXA5 homology arms were synthesized (Genscript). The linearized targeting vector was cotransfected with sgRNA-HOXA5 and Cas9-expressing vectors in 293T cells. 100,000 HOXA5-Snrpn-BFP cells were seeded to 6 well-plates and then mixed with DNMT3A or/and USP11 lentiviral particles in three biological replicates.

### UPS11 depletion using the CRISPR/Cas9 system

Targeting human USP11, 1ug sgRNA (Synthego) (UCUGGGCCCCAGGUUCCUUG for gRNA-1, GCUUCUCCACAAGGAACCUG for gRNA-2), was incubated with 1 μg Cas9 protein (PNA Bio) for 30 minutes at room temperature to obtain Cas9-sgRNA RNPs. Next, the RNPs and 2 × 105 Kelly cells were mixed and electroporated using the optimized electroporation condition of 1,200 V, 20 ms, and two pulses by the Neon transfection system (Thermo FisherScientific).

## Author contributions

T. Y. designed and performed experiments, analyzed the data, and wrote the paper. R. R., V. S., and G. D. assisted with experiments. M. S., and J. R. performed the CRISPR screening. P. G. conceived the project, designed experiments, analyzed the data, and wrote the paper.

## Acknowledgments

We thank the Flow Cytometry Core, the Mass Spectrometry Proteomics Core and the Genetically Engineered Rodent Models Core at Baylor College of Medicine. This work was supported by NIH NCI R01 CA183252 (PG), NIDDK R01 DK092883 (PG), NCI 1P01CA265748-01 (PG), Leukemia & Lymphoma Society Career Development Program (TY), Cancer Research Institute Irvington Postdoctoral Fellow (TY), Japan Society for the Promotion Science Research Fellow (TY), TOYOBO Biotechnology Foundation (TY) and Yamada Science Foundation (TY). I appreciate Dr. Margaret Goodell laboratory members and collaborators.

---

**Supplemental Figure 1.**
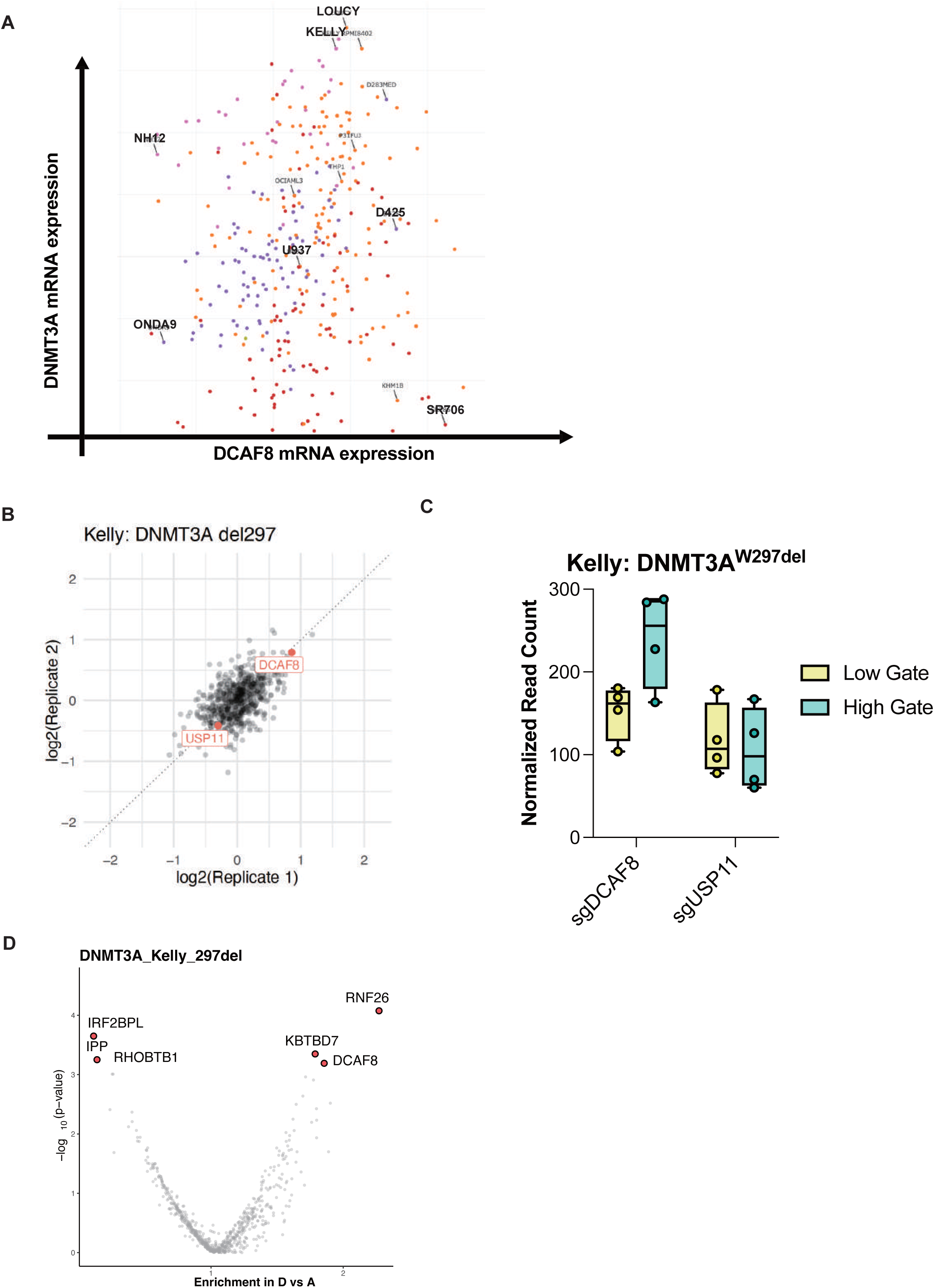
A, The image depicts DNMT3A and DCAF8 mRNA expression in the DepMap (https://depmap.org/portal/) B, The graph depicts the gene enrichment score for DCAF8, USP11 in the CIRSPR screening. C, The graph depicts the read counts for DCAF8, USP11 in the CRISPR screening. D, the graph depicts the gene enrichment score and P value in targeted CRISPR screening for top three factors related to DNMT3A stabilization and destabilization.

**Supplemental Figure 2.**
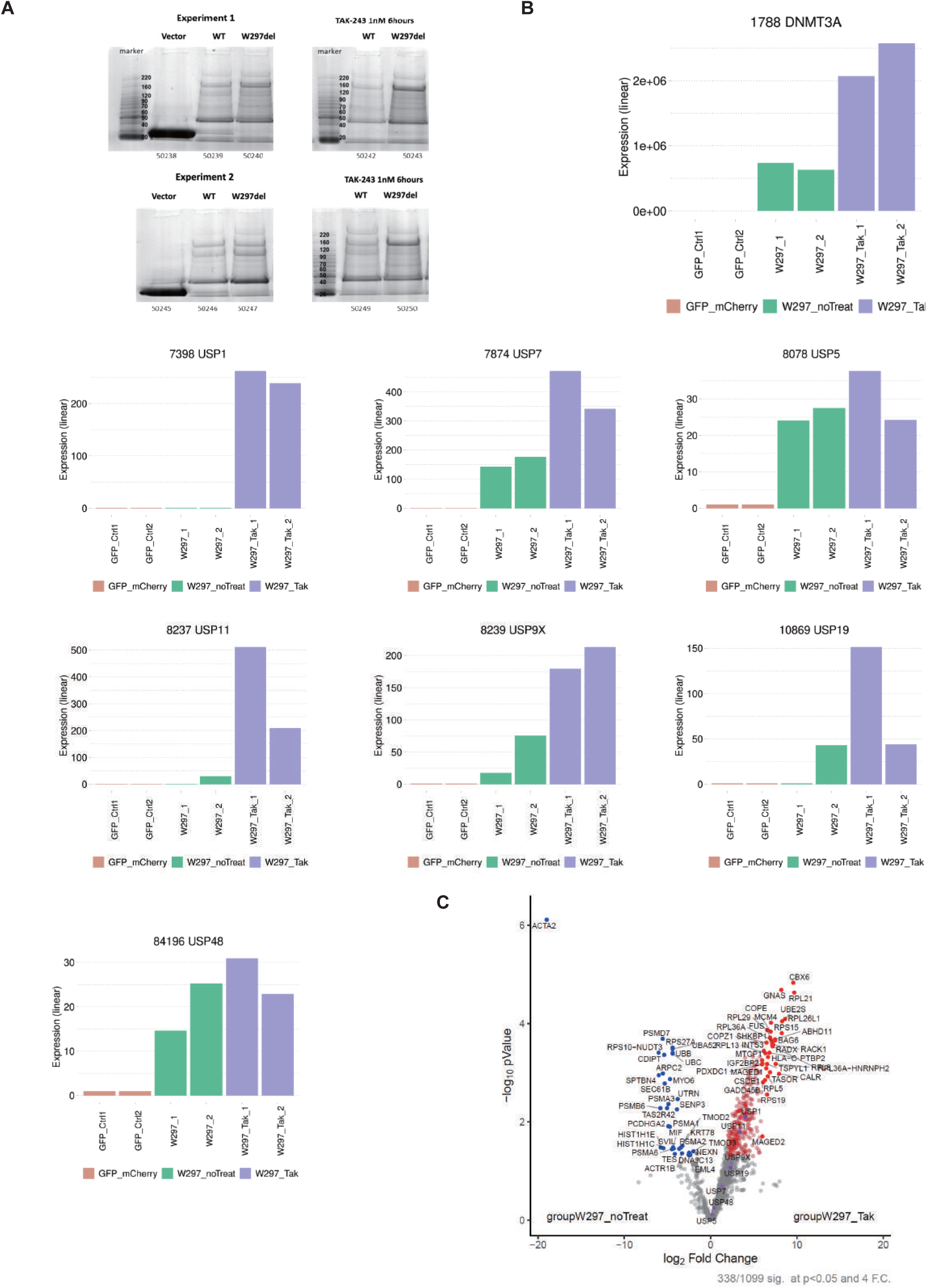
A, The images show representative SDS-PAGE for the DNMT3A IP-MS. B, The graph depicts the protein expression of DNMT3A and deubiquitinating enzymes on IP-MS with DNMT3A. with or without 1nM TAK-243 for 6 hours. C, The graph depicts the enriched score of the interaction between DNMT3AW297del and DNMT3A interacted proteins with or without 1nM TAK-243 for 6 hours.

**Supplemental Figure 3.**
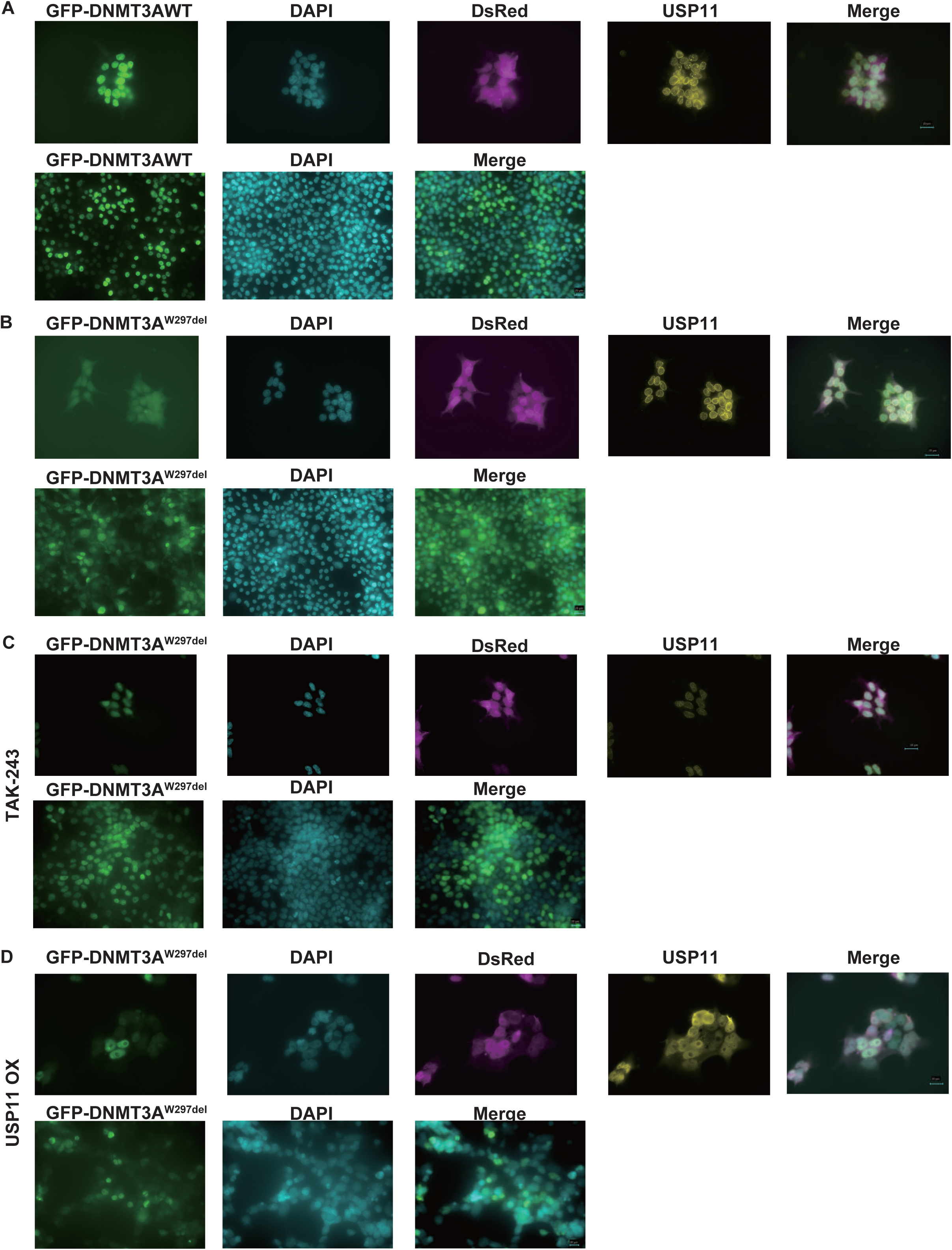
A-D 293T cells were transfected with GFP-DNMT3A WT or-W297del with or without 1nM TAK-243 for 6 hours or USP11 overexpression. We stained anti-USP11 (rabbit) antibodies followed by anti-rabbit Alexa 647 (yellow). DNMT3A expression detected GFP (Green), nuclei and cytoplasm detached, DsRed (Red). Nuclei were visualized with DAPI (Blue). Laser scanning microscopy (Keyence) was used for the imaging.

**Supplemental Figure 4.**
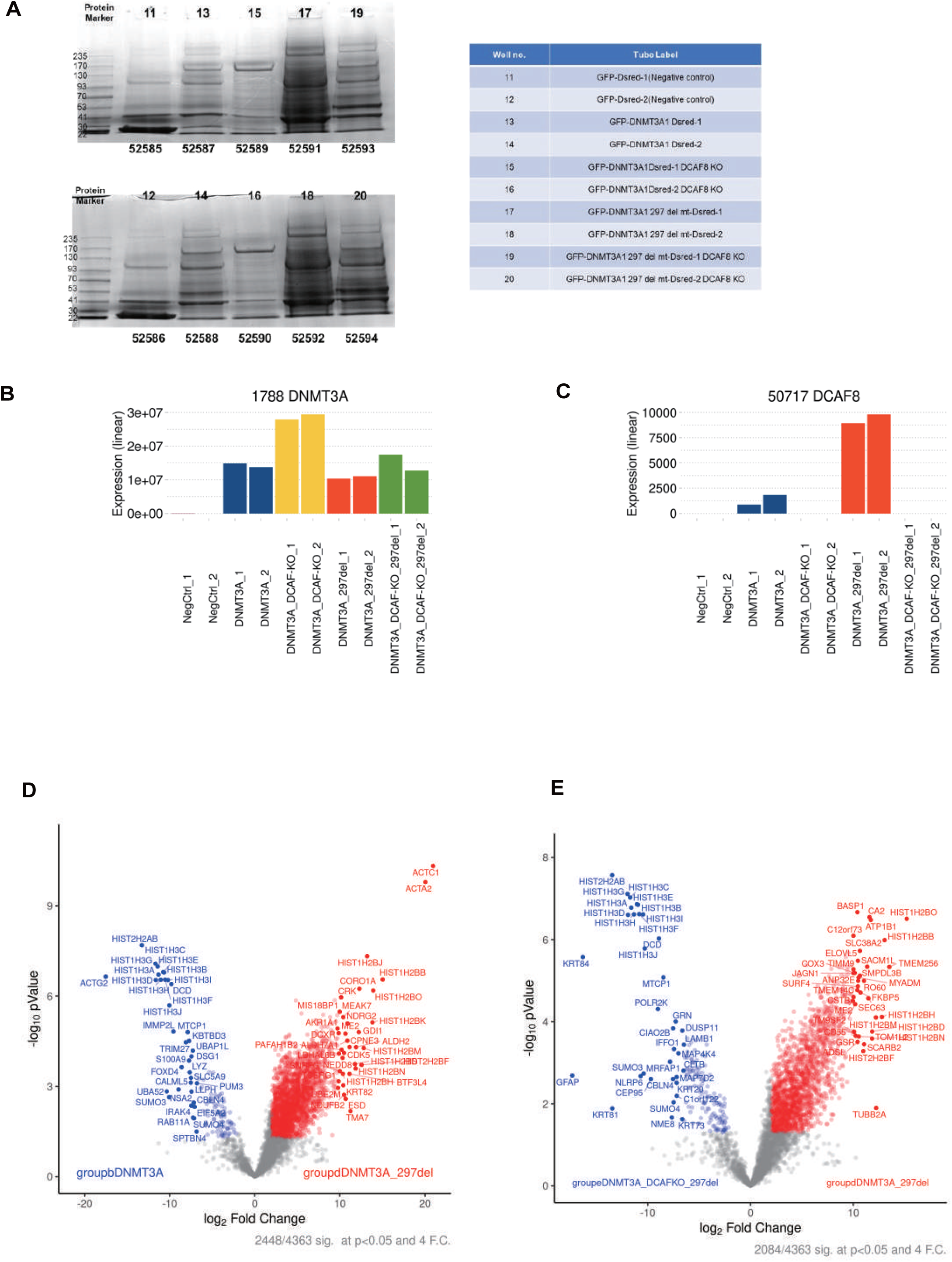
A, The images show representative SDS-PAGE for the DNMT3A IP-MS. B, C, The graph depicts the protein expression of DNMT3A or DCAF8 on IP-MS with DNMT3A. D, E, The graph depicts the enriched score of the interaction between DNMT3A WT and DNMT3AW297del with or without DCAF8.

**Supplemental Figure 5.**
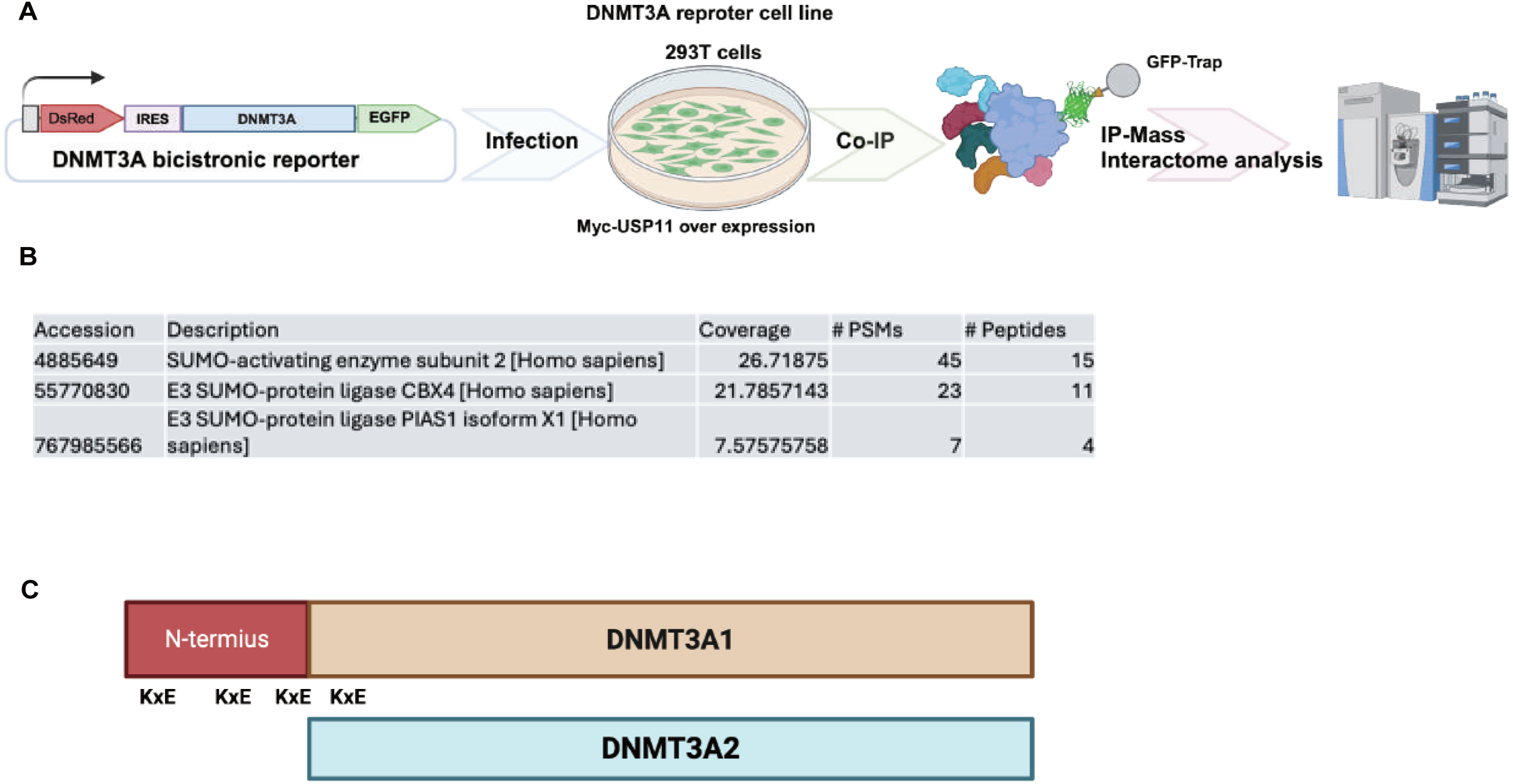
A, Schematic of immunoprecipitation (IP) mass spectrometry to identify DNMT3A E3 SUMO ligases. 293T cells were engineered to constitutively overexpress the indicated bicistronic GFP-DNMT3A WT reporter and transduced Myc-USP11. To subjects cell lysate was performed I P and mass spectrometry analysis. B, The graph depicts the interacted SUMO related proteins with DNMT3A1. C, The graph depicts target protein residues as SUMOylated contain the consensus motif KxE in DNMT3A

**Supplemental Figure 6.**
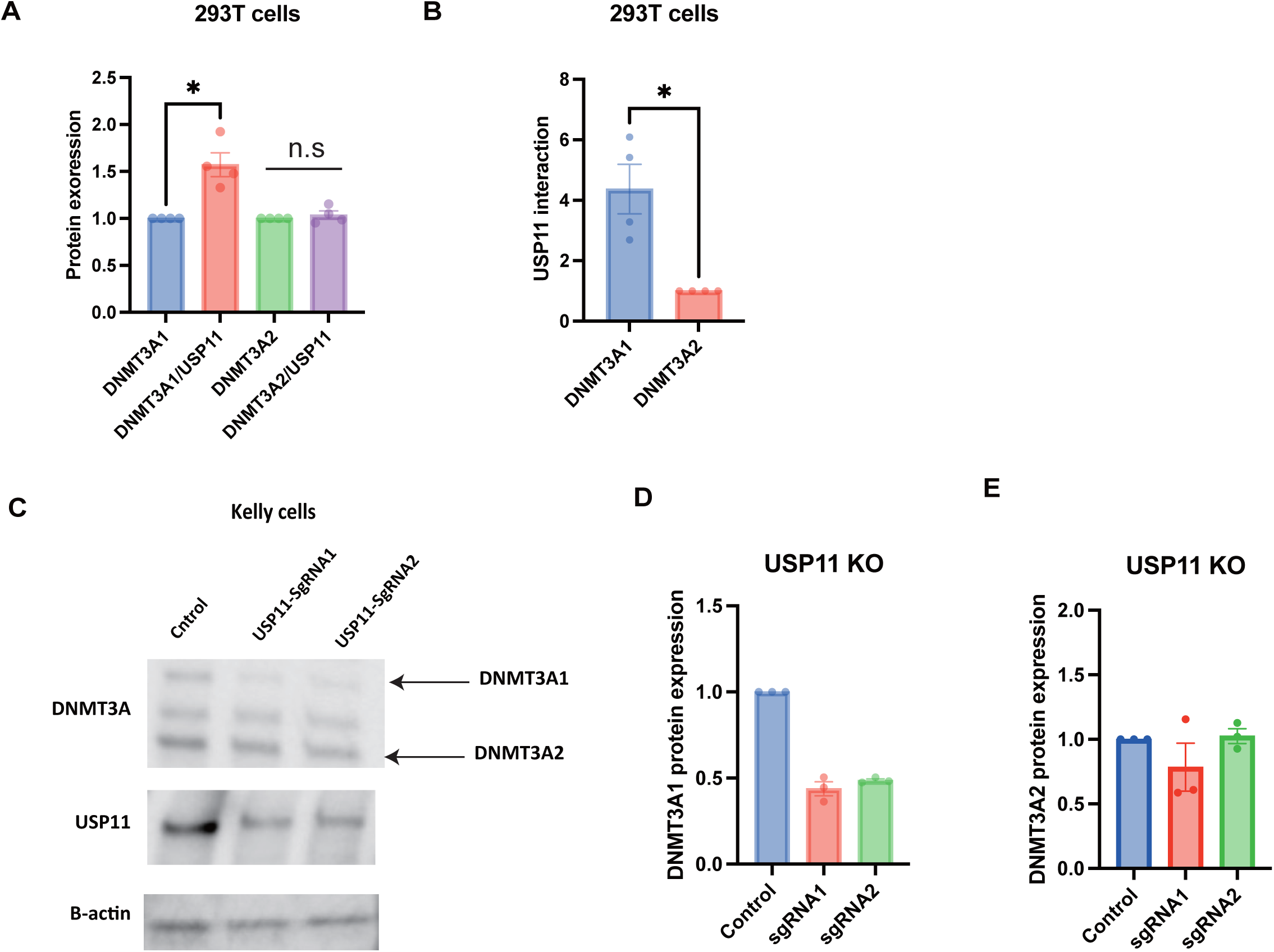
A The graph depicts the protein expression of DNMT3A in input control in the figure 5A. (data are shown as the mean ± S.E.M, *, p = 0.0205, n.s = not significant) Each DNMT3A1 and DNMT3A2 protein expression was set as 1. A The graph depicts the protein expression of Myc-USP11 with FLAG (DNMT3A1 and 2) immunoprecipitation in the figure 5A. (data are shown as the mean ± S.E.M, *, p = 0.0260) C, Kelly cells were transduced with a vector control or two independent gRNAs targeting USP11. The gRNAs showed efficient depletion of USP11. The image depicts DNMT3A, USP11 and B-action protein expression followed by western blotting using DNMT3A, USP11and B-actin antibodies. D, E, The graph depicts the protein expression of USP11, DNMT3A1 and 2 in the supplemental figure 5D.

**Supplemental Figure 7.**
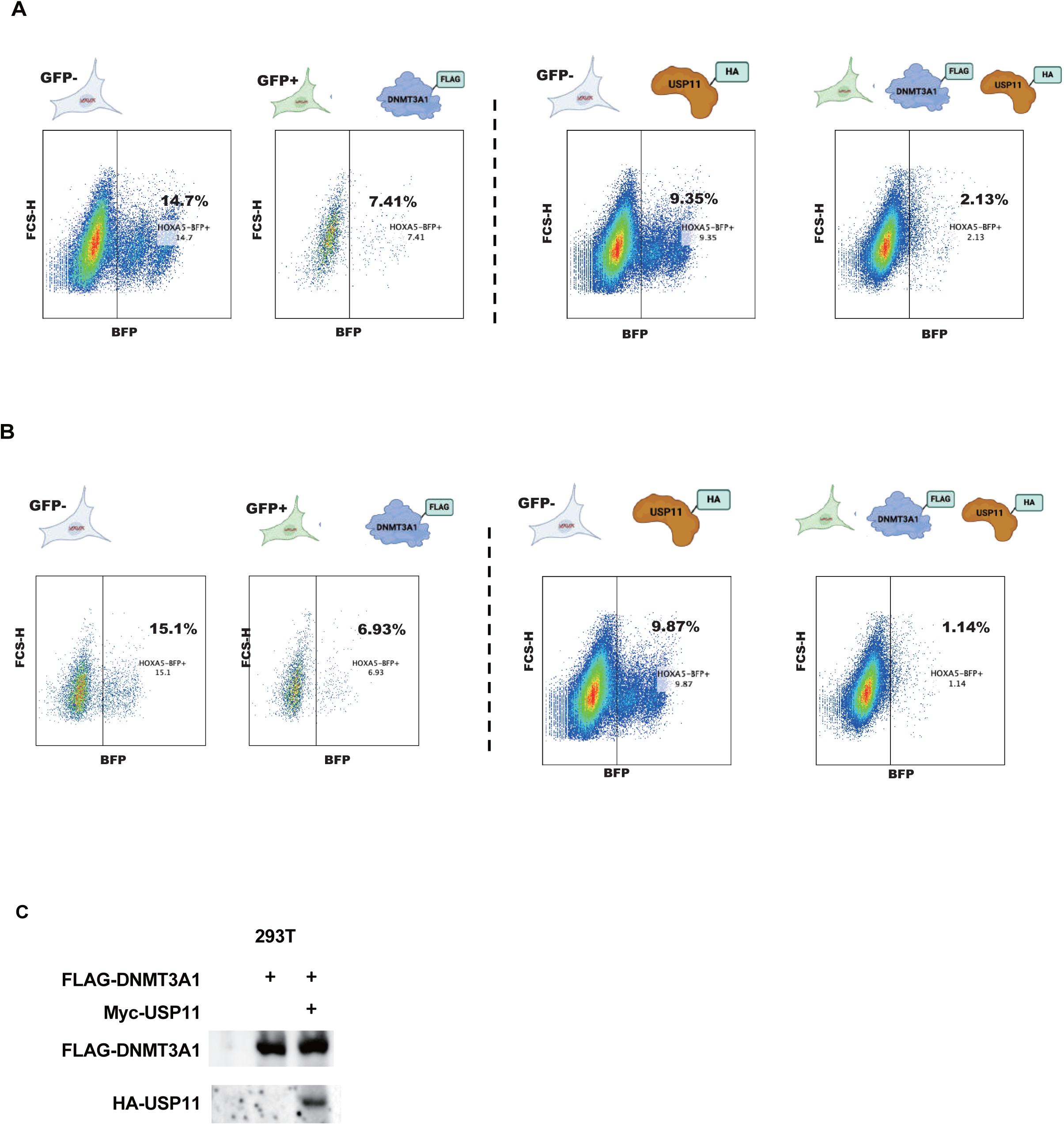
A, B, The figure depicts blue fluorescence intensity in the FLAG-DNMT3A and/or HA-USP11 transduced cells as measured by flow cytometry on day 10. The graphs depicted the BFP positive cells. C, A representative western blot of 293T cells expressing HOXA5-Snrpn-BFP, FLAG-DNMT3A1 and or HA-USP11.

